# Long-read sequencing reveals widespread intragenic structural variants in a recent allopolyploid crop plant

**DOI:** 10.1101/2020.01.27.915470

**Authors:** Harmeet Singh Chawla, HueyTyng Lee, Iulian Gabur, Suriya Tamilselvan-Nattar-Amutha, Christian Obermeier, Sarah V. Schiessl, Jia-Ming Song, Kede Liu, Liang Guo, Isobel A. P. Parkin, Rod J. Snowdon

**Affiliations:** Department of Plant Breeding, Justus Liebig University, Heinrich-Buff-Ring 26-32, 35392 Giessen, Germany; National Key Laboratory of Crop Genetic Improvement, Huazhong Agricultural University, Wuhan, China; Agriculture and Agri-Food Canada, 107 Science Place, Saskatoon, SK S7N OX2, Canada

## Abstract

Genome structural variation (SV) contributes strongly to trait variation in eukaryotic species and may have an even higher functional significance than single nucleotide polymorphism (SNP). In recent years there have been a number of studies associating large, chromosomal scale SV ranging from hundreds of kilobases all the way up to a few megabases to key agronomic traits in plant genomes. However, there have been little or no efforts towards cataloging small (30 to 10,000 bp) to mid-scale (10,000 bp to 30,000 bp) SV and their impact on evolution and adaptation related traits in plants. This might be attributed to complex and highly-duplicated nature of plant genomes, which makes them difficult to assess using high-throughput genome screening methods. Here we describe how long-read sequencing technologies can overcome this problem, revealing a surprisingly high level of widespread, small to mid-scale SV in a major allopolyploid crop species, *Brassica napus*. We found that up to 10% of all genes were affected by small to mid-scale SV events. Nearly half of these SV events ranged between 100 bp to 1000 bp, which makes them challenging to detect using short read Illumina sequencing. Examples demonstrating the contribution of such SV towards eco-geographical adaptation and disease resistance in oilseed rape suggest that revisiting complex plant genomes using medium-coverage, long-read sequencing might reveal unexpected levels of functional gene variation, with major implications for trait regulation and crop improvement.

## Introduction

The recent allopolyploid species *Brassica napus* L. (oilseed rape/canola/kale/rutabaga; genome AACC, 2n=38) evolved rapidly into a globally important crop. Genome assembly and resequencing of *B. napus* (Chalhoub et al. 2014) revealed a highly complex and strongly duplicated genome with an unexpected extent of segmental exchanges among homoeologous chromosomes. In synthetic *B. napus* accessions, genome structural variants frequently span whole chromosomes or chromosome arms (Chalhoub et al. 2014, Samans et al. 2017). Naturally formed *B. napus* also shows widespread homoeologous exchanges, with similar distribution patterns (Hurgobin et al., 2018; Samans et al., 2017), that apparently arose during the allopolyploidisation process (Leflon et al., 2006; Nicolas et al., 2007; Szadkowski et al., 2010). The wide extent of segmental deletion/duplication events in both synthetic and natural *B. napus* has been confirmed using other genome-wide analysis methods, for example visualization based on mRNAseq data (He et al., 2017) or deletion calling from SNP array data (Gabur et al., 2018; Grandke et al., 2016). Critically, numerous examples have connected genome SV in *B. napus* to important agronomic traits (Gabur et al., 2018; Gabur et al., 2019; Liu et al., 2012; Stein et al., 2017). These studies revealed the important role of SV in the creation of *de novo* variation for adaptation and breeding, however the methods used were not yet capable of resolving SV at gene scale.

A first example of intragenic SV impacting quantitatively inherited traits in *B. napus* was reported by Qian et al. (2016), who demonstrated that deletion of exons 2 and 3 from a *B. napus* orthologue of Mendel’s “Green Cotyledon” gene (the Staygreen gene *NON-YELLOWING 1; NYE1*) associated with quantitative variation for chlorophyll and oil content. Unfortunately, such small deletions are challenging to reliably detect using short-read sequencing or low-cost marker arrays, so that their genome-wide extent could not yet be investigated in detail. In this study, using *B. napus* as an example for a plant genome with widespread structural variation, we demonstrate the power of whole-genome long-read sequencing for high-resolution detection of intragenic SV. The results reveal widespread functional variation on a completely unexpected scale, suggesting that small to mid-scale SV may be a major driver of functional gene diversity in this recent polyploid crop. With the growing accessibility, accuracy and cost-effectiveness of long-read sequencing, our results suggest that there could be enormous promise in revisiting complex crop genomes to discover potentially novel functional SV which has previously been overlooked.

## Results and discussions

### Long read sequencing reveal novel SV diversity in *B. napus*

We sequenced 4 *B. napus* accessions with long reads using the Oxford Nanopore Technology (ONT) an 8 further accessions using the Pacific Biosciences (PacBio) platform (obtained from Song et al. (2020)). The genotype panel included three vernalisation-dependent winter-type accessions, 3 vernalisation-independent spring-type accessions, 4 semi-winter accessions and 2 synthetic *B. napus* accessions (a winter-type and a spring-type). All accessions were sequenced to between ~30x and ~50x whole-genome coverage (between 30 and 50 Gb of data). Reads were aligned to the *B. napus Darmor-bzh* version 4.1 reference genome (Chalhoub et al., 2014) using the long-read aligner NGMLR (https://github.com/philres/ngmlr) (Sedlazeck et al., 2018) and called for genome-wide SV using the SV-calling algorithm Sniffles (Sedlazeck et al., 2018). N50 values ranging from 10,552 to 15,369 bp were obtained for the 8 PacBio datasets, while in the 4 ONT datasets the N50 ranged from 10,756 to 28,916 bp (Table 1, Supplementary Table S1). After aligning to the *Darmor-bzh* v4.1 reference genome, the total number of SV events called by Sniffles ranged from 51,463 to 108,335. To minimise false-positive calls derived from reference mis-assemblies, we followed a highly stringent quality-filtering approach (details in Supplementary Materials) that removed 54.4-59.4% of the total predicted SV. This procedure resulted in a final set of 27,107 to 44,516 high-quality SV events (Table 1). To evaluate the impact of assembly errors on SV calling rates, we compared results after aligning (using the same procedure) to a pseudo-reference constructed by combining the high-quality long-read reference assemblies of *Brassica rapa* (A subgenome) and *Brassica oleracea* (C subgenome) published recently by Belser et al. (2018). Using this pseudo-reference assembly we detected between 41,436 and 50,907 quality filtered SV across the 12 *B. napus* genotypes. There are two possible explanations for the higher number of SV. Firstly, the pseudo-reference assembly (957 Mbp) is nearly 10 percent larger than the *B. napus* Darmor-*bzh* v4.1 reference (849.7 Mbp). Secondly, SV detected using the pseudo-reference assembly will also reflect genomic differences between the unknown diploid progenitors of *B. napus* and the two diploid genotypes from which this pseudo-assembly was generated. To further validate our SV detection approach, we therefore compared the number of SV per megabase, detected using the two different genome assemblies for each of the 19 chromosomes across 12 genotypes. This showed a correspondence of 77.08 percent, suggesting that the latter may be the predominant cause.

**Table 1:** Number and size distributions of SV detected in 12 *B. napus* genotypes. ONT: Oxford Nanopore Techhnologies; PacBio: Pacific Biosciences; SV: Structural variant.

| Genotype | Data type | Ecotype | N50 for raw reads | Quality filtered SV | Intra-genic SV | Minimum size of SV | Maximum size of SV | Median size SV |
| --- | --- | --- | --- | --- | --- | --- | --- | --- |
| **Express 617** | ONT | Winter | 10,756 | 27,107 | 5,383 | 31 | 16,931 | 341 |
| **Quinta** | Pacbio | Winter | 14,192 | 32,349 | 7,286 | 31 | 15,869 | 353 |
| **Tapidor** | Pacbio | Winter | 14,448 | 32,757 | 7,479 | 31 | 15,289 | 344 |
| **ZS11** | Pacbio | Semi-winter | 10,552 | 37,496 | 9,165 | 31 | 11,312 | 281 |
| **Zheyu7** | Pacbio | Semi-winter | 12,370 | 38,590 | 9,226 | 31 | 17,001 | 305 |
| **Gangan** | Pacbio | Semi-winter | 14,064 | 35,560 | 8,542 | 31 | 14,264 | 335 |
| **Shengli** | Pacbio | Semi-winter | 13,828 | 39,622 | 9,697 | 31 | 12,207 | 321 |
| **PAK85912** | ONT | Spring | 28,916 | 23,177 | 5,172 | 31 | 28,777 | 584 |
| **N99** | ONT | Spring | 27,139 | 34,848 | 7,700 | 31 | 26,183 | 509 |
| **Westar** | Pacbio | Spring | 13,810 | 37,138 | 8,769 | 31 | 17,615 | 332 |
| **R53** | ONT | Winter<br>synthetic | 11,253 | 33,851 | 7,929 | 31 | 12,635 | 296 |
| **No2127** | Pacbio | Spring<br>synthetic | 15,369 | 44,516 | 10,869 | 31 | 15,565 | 304 |

After alignment to the Darmor-*bzh* v4.1 reference genome, the median detected SV size across the 12 accessions ranged from 296 bp to 584 bp. The spring-type accessions N99 and PAK85912 had the largest median SV size (509 and 584 bp, respectively), which might be attributable to the longer read lengths for these two genotypes (N50 = 27,139 bp and 28,916 bp, respectively) (Figure 1A). The largest SV event (34,848 bp) was also detected in the spring-type accession N99, suggesting that read length plays a critical role in the ability to detect large and complex SV events. Around half of all detected, high-confidence SV events (46.8 to 53.2 % across the 12 genotypes) ranged in size from ~100-1000 bp (Supplementary Table S2). These small SV represent a novel genetic diversity resource that was previously unnoticed due to the insufficient resolution of high-throughput genotyping platforms such as SNP genotyping arrays and a very high false-positive rates (up to 89%) of short-read sequencing data (Mahmoud et al., 2019; Sedlazeck et al., 2018).

**Figure 1:**
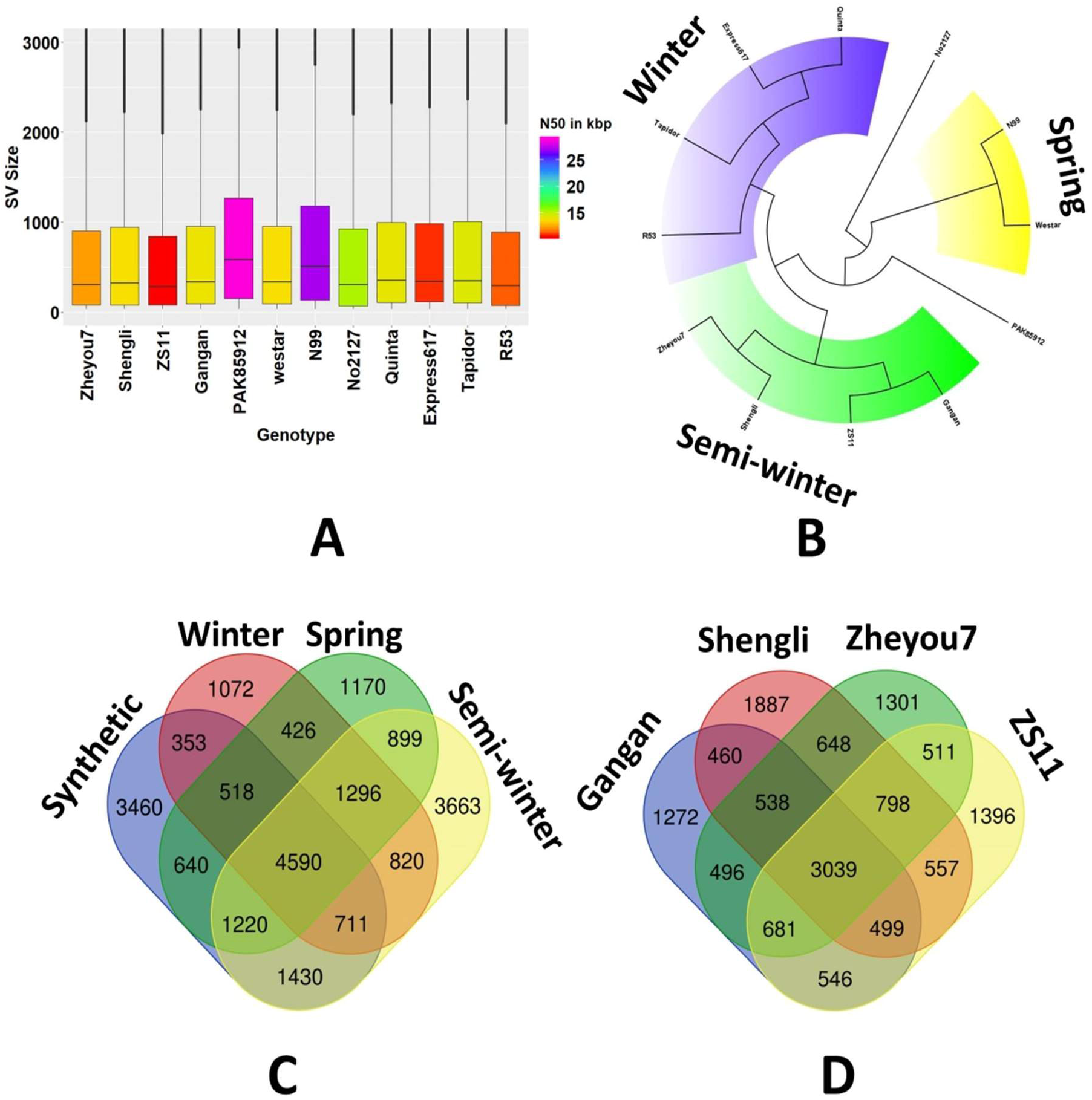
**A.** Box plots showing size distributions of SV events detected in 12 B.napus genotypes. **B.** Maximum likelihood tree showing genetic relationships among 12 B. napus genotypes based solely on genome-wide SV events, revealing clear clustering into the appropriate ecogeographical morphotype groups. C. Venn diagram showing the numbers of common or unique genes carrying intragenic SV events across three divergent ecotypes and synthetic B. napus, respectively. **D.** Venn diagram representing the numbers of common or unique genes carrying intragenic SV events across 4 semiwinter *B. napus* accessions.

### Subgenomic differences in SV frequency

Comparison of subgenomic SV frequency revealed significantly higher numbers of small-to mid-scale SV per megabase in the *B. napus* A subgenome than the C subgenome in all twelve analysed genotypes (Figure 6 A and B, Supplementary Table S4). This reflects a corresponding subgenomic bias also observed for large-scale SV in *B. napus* (Samans et al. 2017), this could also be attributable to repeated introgressions from the A genome of *B. rapa* during the breeding history of *B. napus* (Lu et al., 2019). Samans et al. (2017) reported a significant enrichment for large-scale segmental deletions in the C-subgenome of *B. napus* resulting from homoeologous exchanges. In contrast, we observed no bias for small to mid-scale deletions in the C-subgenome of the 12 sequenced *B. napus* accessions (Supplementary table S6). This indicates that a different molecular mechanism may be responsible for the generation of large and small to mid-scale SV events in the rapeseed genome. Unexpectedly, we found that between 5% (Express 617) and 10% (No2127) of all genes detected in the twelve accessions were affected by small to mid-scale SV events. This represents a previously completely unknown extent of functional gene modification as a result of post-polyploidisation genome restructuring. It also underlines the massive selection potential arising from intergenomic disruption during the act of allopolyploidisation (Nicolas et al., 2007; Nicolas et al., 2008; Szadkowski et al., 2010), and the great significance of post-polyploidisation intergenomic restructuring for polyploid crop evolution (Samans et al., 2017).

### Small to mid-scale SV underlining eco-geographical differentiation in *B. napus*

As expected, strong SV differentiation from the winter-type oilseed reference genotype Darmor-*bzh* was found in the divergent semi-winter and spring ecotypes, and in genetically distant synthetic *B. napus* accessions R53 and No2127 (Figure 2). Unexpectedly, however, the winter-type accessions Express 617, Tapidor and Quinta also showed high levels of SV compared to Darmor-*bzh*, despite a related breeding history and partially shared pedigree (e.g. Express 617). According to (Lu et al., 2019), who used whole-genome resequencing data to investigate the species origin and evolution of *B. napus*, spring and semi-winter types arose only very recently (<500 years) from winter-types. Our data concur with this assumption, with fewer genes carrying SV in winter-type accessions (1072) than in spring (1170) or semi-winter (3663) ecotypes (Figure 1C). Furthermore, we also detected small to mid-scale SV within each ecotype, for example 1272-1887 genes carrying unique SV events were found among the four semi-winter accessions (Figure 1D). The unexpectedly high structural gene diversification both between and within ecotypes suggests that *de novo* generation of small to mid-scale SV may also be ongoing in recent breeding history. Overall, 4590 of the called intragenic SV were common among the four *B. napus* forms, indicating putative SV events specific to Darmor-*bzh*. These could possibly be attributed to errors in the Darmor-*bzh* reference assembly, however the similar number of unique intragenic SV detected only in semi-winter types (3663) suggests that this frequency is not unexpected in the context of the other results. Repeating the analysis with the concatenated pseudo-reference from *B. rapa* plus *B. oleracea* gave comparable results (6248 common among all sequenced *B. napus* forms, 2919 unique to semi-winter ecotypes).

**Figure 2:**
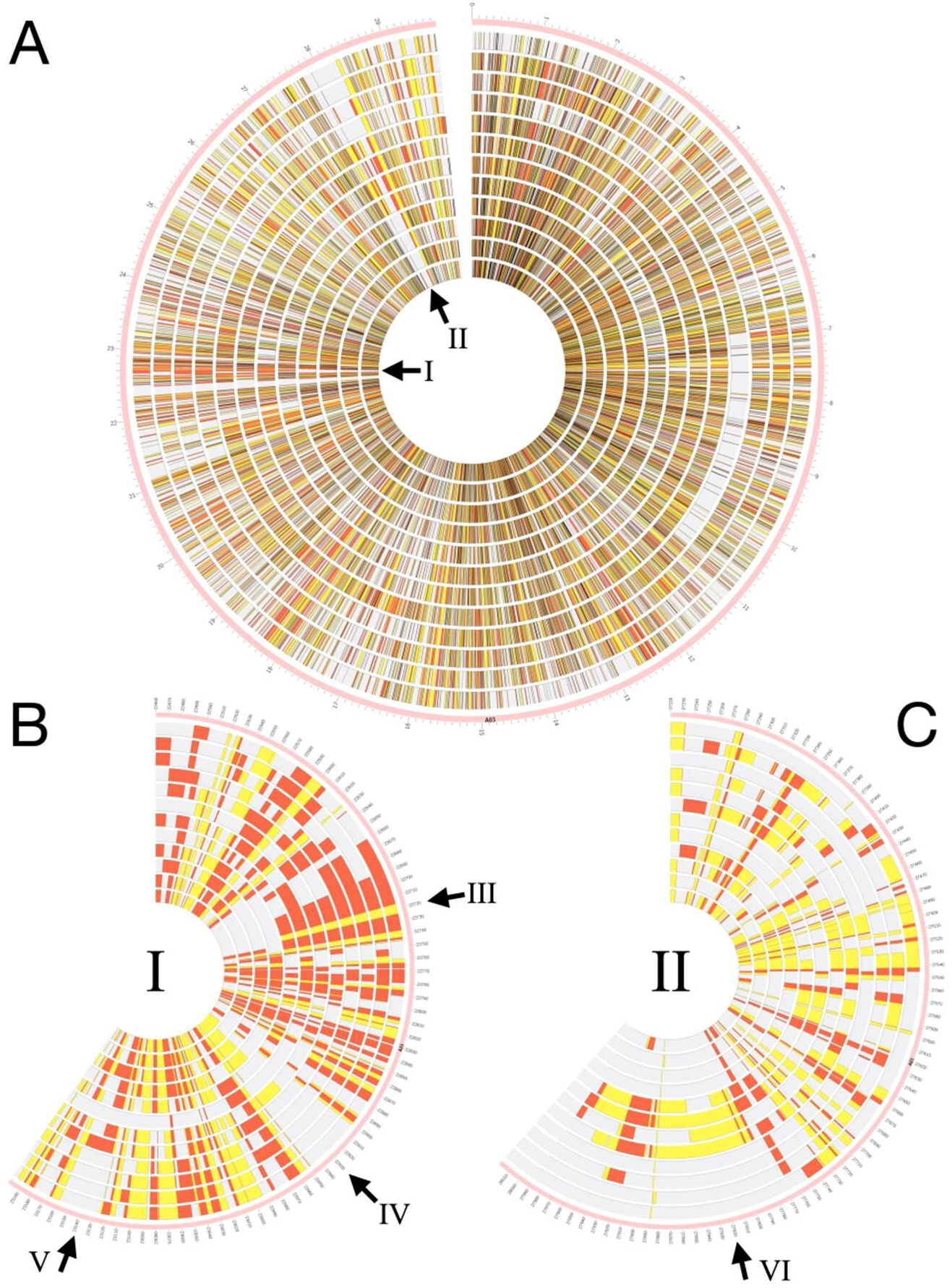
**A:** Circos plot showing small to mid-scale insertion and deletion events in 12 *B. napus* accessions, using chromosome AO3 as an example. Each track represents a single accession in the following order from outside to inside: Express 617, Quinta, Tapidor, R53 (all winter-type), No2l27, N99, Westar, PAK85912 (spring-type), Gangan, Shengli, Zheyou7 and ZS11 (semi-winter type). Deletions are represented by yellow blocks, whereas insertions are shown by red blocks. Darker blocks in (A) represent regions containing both deletions and insertions in different genotypes. Arrows I and II mark selected segmental SV events specific for a particular ecotype. B: Expanded view of the chromosome segment depicted by arrow I in A. Arrow III represents a 50 kbp region containing segmental deletion and insertion events detected in all winter and spring ecotypes but not in the semiwinter-types. Arrow IV indicates a 40 kbp region containing segmental deletions detected only in the four semi-winter types and three of spring-types. Arrow V indicates a 40 kbp region containing segmental insertions detected only in the four semi-winter types and one of the spring-types. C. Expanded view of the chromosome segment depicted by arrow II in A. Arrow VI indicates a 120 kbp region containing segmental insertions only in the four spring-types

To evaluate the influence of SV on eco-geographical adaptation and potential species diversification, we constructed a maximum likelihood (ML) tree for the 12 *B. napus* lines based solely on SV detected using long read sequencing data. The resulting tree (Figure 1B) comprised 3 divergent clades representing 3 ecotypes of *B. napus* (winter, semi-winter and spring). In contrast to genetic clustering based on genome-wide SNP data, which reveals high sequence diversification between synthetic and natural *B. napus* (Bus et al., 2011), the two synthetic accessions R53 and No2127 did not fall into separate clades. Instead, the winter-type R53 clustered closest together with the natural winter-type accessions and the spring-type No2127 clustered with the natural spring-type accessions. This suggests that small to mid-size SV events originating during or immediately after allopolyploidisation might rapidly confer ecogeographical adaptation. Although hundreds to thousands of genes carrying unique SV events were detected in each individual accession, the intriguing observation that their cumulative clustering reflects ecogeographical adaptation forms suggests a possible key role of SV in rapid functional adaptation. Overall, the distribution and frequency of SV events in all investigated accessions suggest that small to mid-scale SV may be a major, previously unknown source of functional genetic variation in *B. napus*.

Unfortunately, a catalogued and validated “truth set” of genomic SV is not yet established for *B. napus* or other complex plant genomes. This makes it crucial to validate SV predicted from long reads using independent validation methods. On the other hand, manual verification of thousands of SV events (for example using PCR) is not realistic. To obtain first insight into the validity of the SV called using our pipeline, we selected relevant, potentially functional examples representing possible functional mutations in flowering-time and disease resistance-related genes. We validated the detected SV events using different independent methods in a total of 4 *B. napus* genotypes including two springs, one winter and a synthetic.

### Small to mid-scale SV events impact *B. napus* flowering time pathway genes

In order to understand the impact of gene scale re-arrangements on eco-geographical adaptations in *B. napus*, we examined the abundance of SV in the known *B. napus* orthologs of all known genes from the *Arabidopsis* flowering-time pathway. Whereas most of these genes are present in only a single copy in *Arabidopsis*, all have multiple duplicates in *B. napus* (Schiessl et al., 2014). Although many *B. napus* flowering-time gene orthologues are known to be affected by copy-number variation, the exact positions of copy-number variants and other small to mid-scale forms of SV could not be determined from previous, short-read resequencing data (Schiessl et al., 2017). Using long-read data, we found that 44 of 178 flowering-time pathway genes, including numerous key regulatory genes, contain one or more small to mid-scale insertions or deletions. For example, we detected a 90 bp insertion in an orthologue of *Vernalisation Insensitive 3* on chromosome C03 (*BnVIN3.C03, BnaCO3gl298OD*) in 3 out of 12 total genotypes, Express 617, No2127 and Zheyou7 (Figure 3A). Successful validation of this insertion via PCR, using primers designed from the SV-flanking sequences, is shown in Figure 3B. The same insertion was undetectable using only the short read sequence-capture data of Schiessl et al. (2017). In two out of three spring accessions, N99 and PAK85912, we detected a 2.8 kbp insertion in a *B. napus* orthologue of the key vernalisation regulator *Flowering Locus C* (*BnFLC.A02, BnaA02g00370D*) a variant previously reported by Chen et al. (2018) to be causal for early flowering.

**Figure 3:**
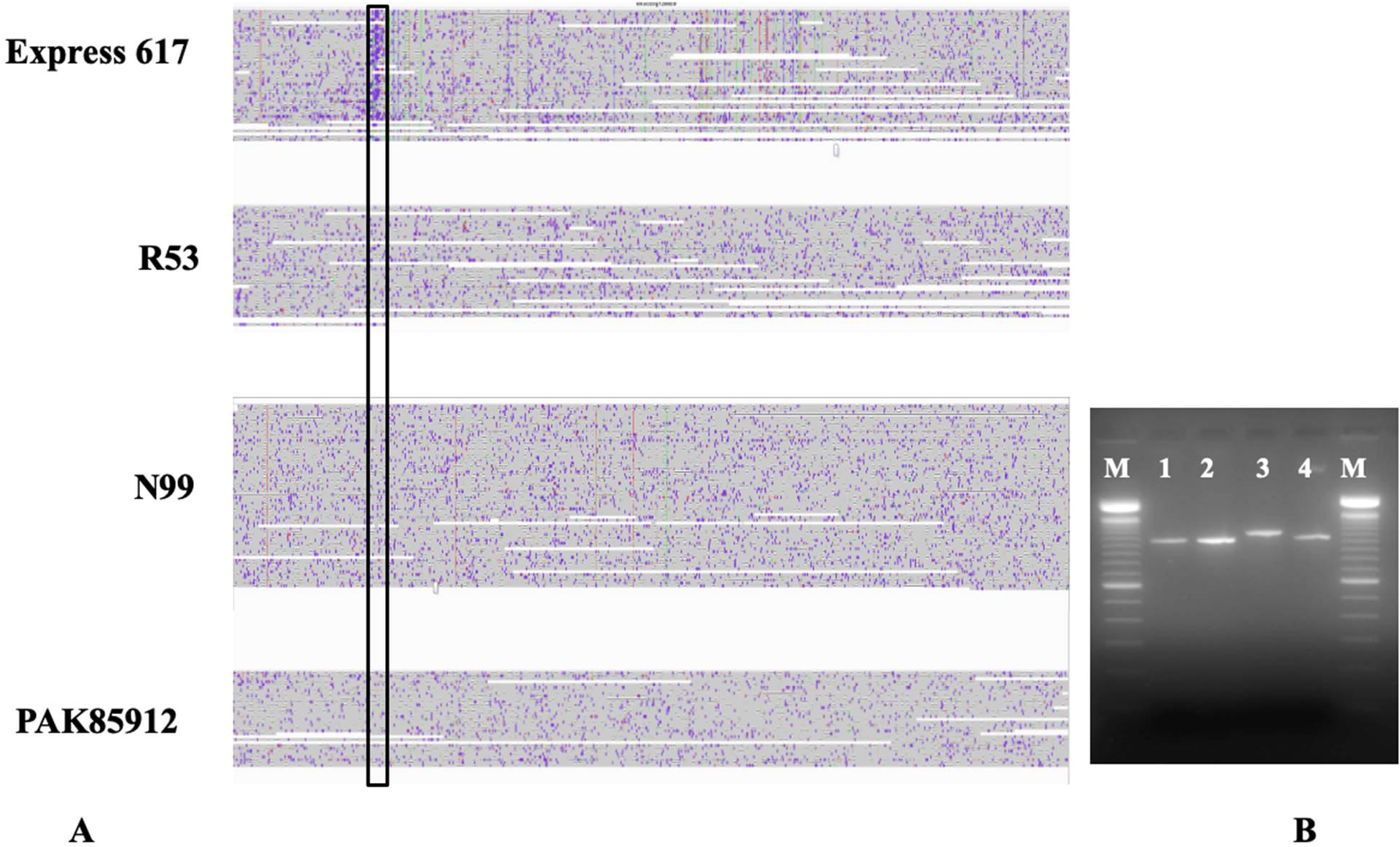
**A.** 90 bp insertion (highlighted in the black box) in an orthologue of *Vernalisation Insensitive 3* on chromosome C03 *Ų3nVIN3.CO3*) revealed by aligning ONT reads from 4 different genotypes to the Darmor-6∑7z reference version 4.1 (detected only in Express 617). B. Agarose gel image of PCR product from the same insertion. M represents a 100 bp ladder and 1-4 represent PCR product originating from N99, PAK85912, Express 617 and R53 respectively. As expected Express 617 exhibits a PCR product size of 1090 bp whereas a 1000 bp product is observed for the rest of three genotypes.

In a second case study, we analysed SV events in key vernalisation genes that differentiate between the vernalisation-dependent and vernalisation-independent *B. napus* accessions in our panel. A number of interesting, putative functional variants were detected. For example, we detected a 288 bp deletion (Figure 5) in all the spring and semi-winter accessions (except for ZS11) in *BnFT.A02 (BnaA02g12130D*). This *FT* ortholog on chromosome A02 has been reported to be significantly associated with flowering-time variation in a worldwide collection of rapeseed accessions (Wu et al., 2019). *BnFT.A02* was also found to be differentially expressed among winter, spring and semi-winter type *B. napus* by Wu et al.

(2019), therefore we scanned for SV in the putative promoter region for this gene. We identified a 1.3 kbp deletion between 6,365,143 and 6,366,504 bp on chromosome A02, exclusively present in all 4 spring accessions, which was situated approximately 10kbp upstream from the start codon of *BnFT.A02* (Figure 5).

### Intragenic SV events associate with disease resistance in oilseed rape

Samans et al. (2017) and Hurgobin et al. (2018) revealed that defence-related R-genes involved in monogenic resistance are particularly enriched in genome regions affected by large-scale SV in *B. napus*. In a third case study related to a prominent disease resistance in oilseed rape, we investigated the impact of SV in resistance-related genes co-localising with QTL for quantitative disease resistance in a bi-parental cross between the sequenced accessions Express 617 and R53. These two accessions differ strongly in their resistance reaction to the important fungal pathogen *Verticillium longisporum* (Obermeier et al., 2013), and SV detected between the two parental lines were selected for validation based on their co-localization to resistance-related genes in corresponding resistance QTL (see Supplementary Methods for selection criteria for PCR validation of SV events). Most interestingly, we identified a 700 bp deletion in R53 that caused the loss of three exons of a *4-Coumarate:CoA Ligase* (*4CL*) gene (*BnaC05g15830D*). In the genetic map from the Express 617 x R53 mapping population, this gene is located within a major QTL for *V. longisporum* resistance on *B. napus* chromosome C05 (Obermeier et al., 2013). 4CL is a critical enzyme involved in the phenylpropanoid pathway (Li et al., 2015) and Obermeier et al. (2013) reported that major QTL for phenylpropanoid compounds co-localized with the QTL for *V. longisporum* resistance in the Express 617 x R53 mapping population. Locus-specific PCR primers, spanning the putative SV predicted by the long sequence reads, amplified 900 bp and 200 bp fragments for Express 617 and R53, respectively (Figure 4 A and B), confirming the expected 700 bp deletion. Re-screening of the PCR markers for the 700 bp deletion in the doubled haploid mapping population from Express 617 x R53 confirmed their co-localisation with the QTL and a strong effect on resistance of up to R^2^= 19.4%.

**Figure 4:**
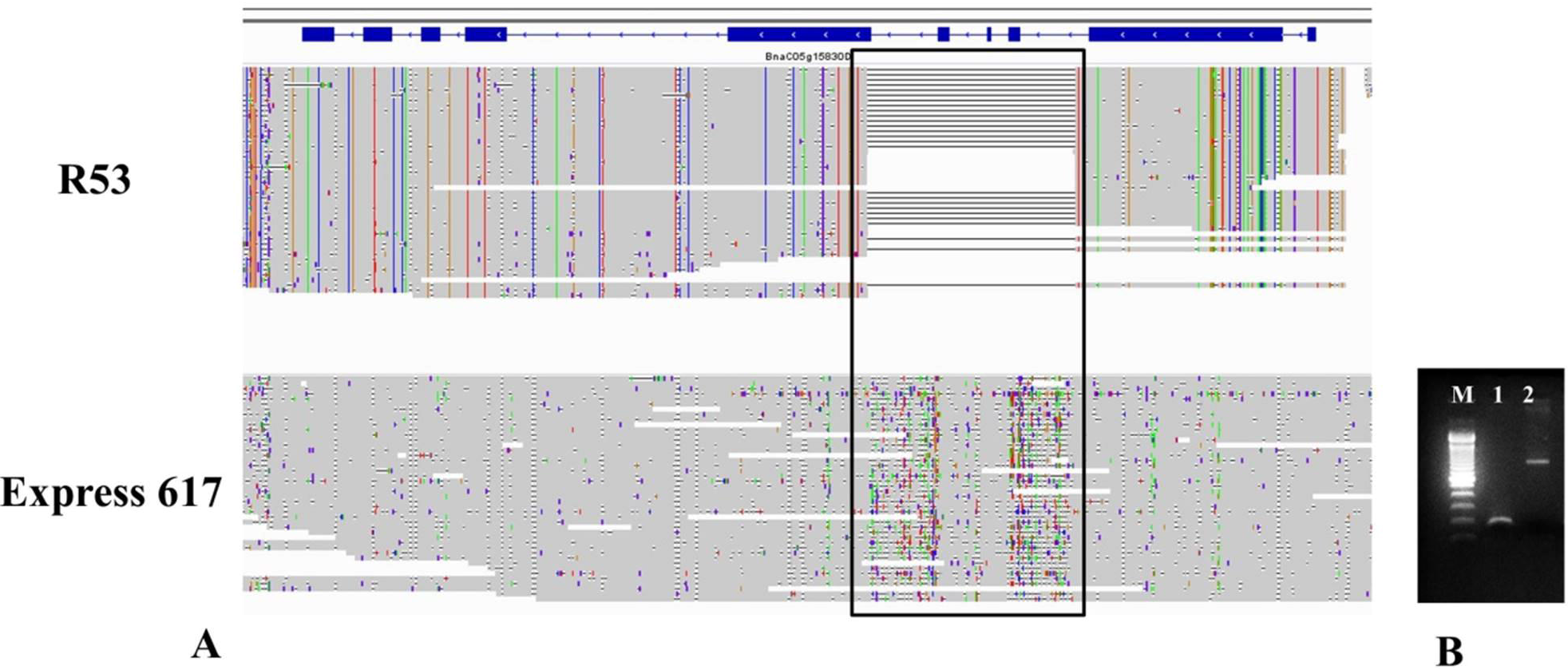
**A.** 700 bp deletion (highlighted in the black box) in R53 that caused the loss of three exons of a *4-Coumarate: Co A Ligase (4CL*) gene (*BnaCO5gl 583OD*)**. B.** Agarose gel image of PCR product from the same deletion. M represents a 100 bp ladder and 1,2 represent PCR product originating from R53 and Express 617 respectively. As expected Express 617 exhibits a PCR product size of 900 bp whereas R53 shows a band at 200 bp

**Figure 5:**
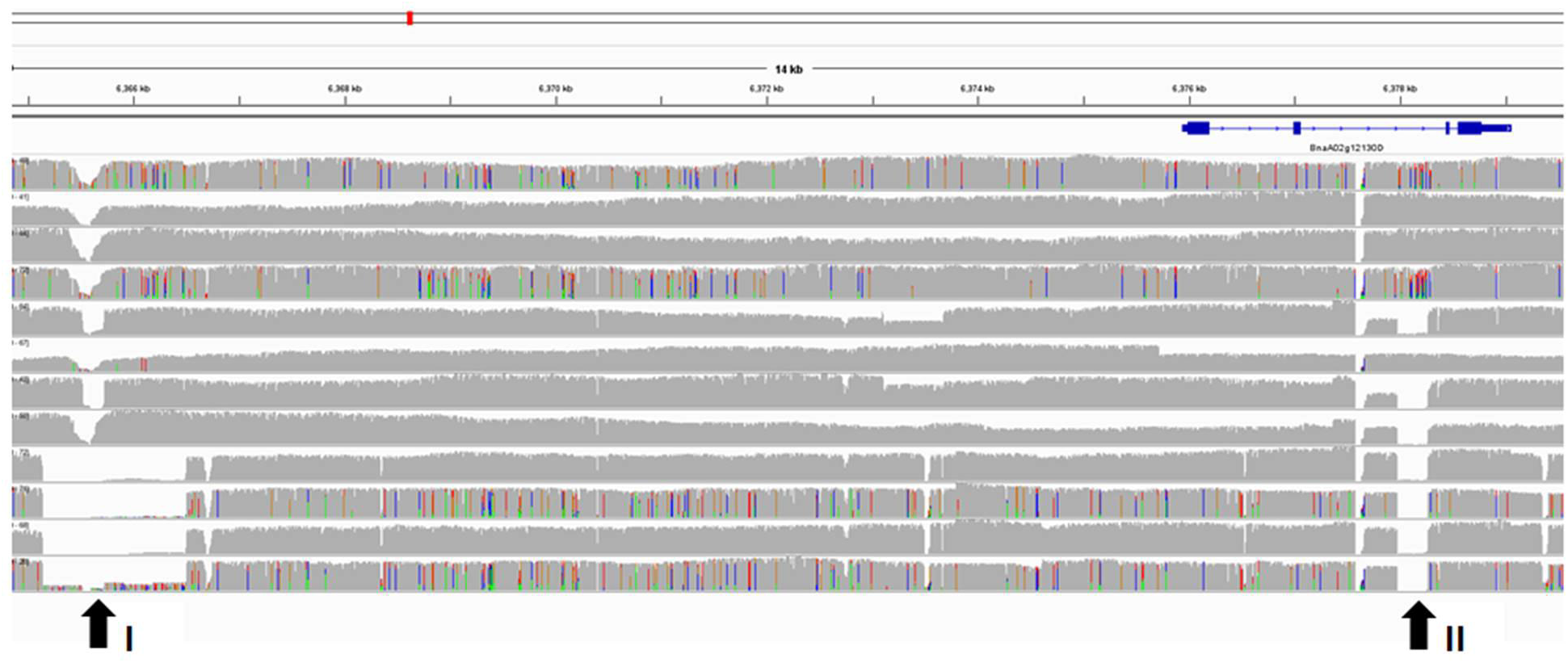
Each track represents a single genotype in the following order from top to bottom: Express 617, Tapidor, Quinta, R53, Shengli, ZS11, Gangan, Zheyou7, No2127, N99, Westar and PAK85912. Arrow I indicate a 1.3 kbp deletion in putative promoter region in *B⊓FT.AO2* (*BnaAO2gl213OD*) for all 4 spring accessions (No2127, N99, Westar and PAK85912). Arrow II indicates a 288 bp deletion in all the spring and semi-winter accessions (except for ZS11) in *B⊓FT.AO2*.

**Figure 6:**
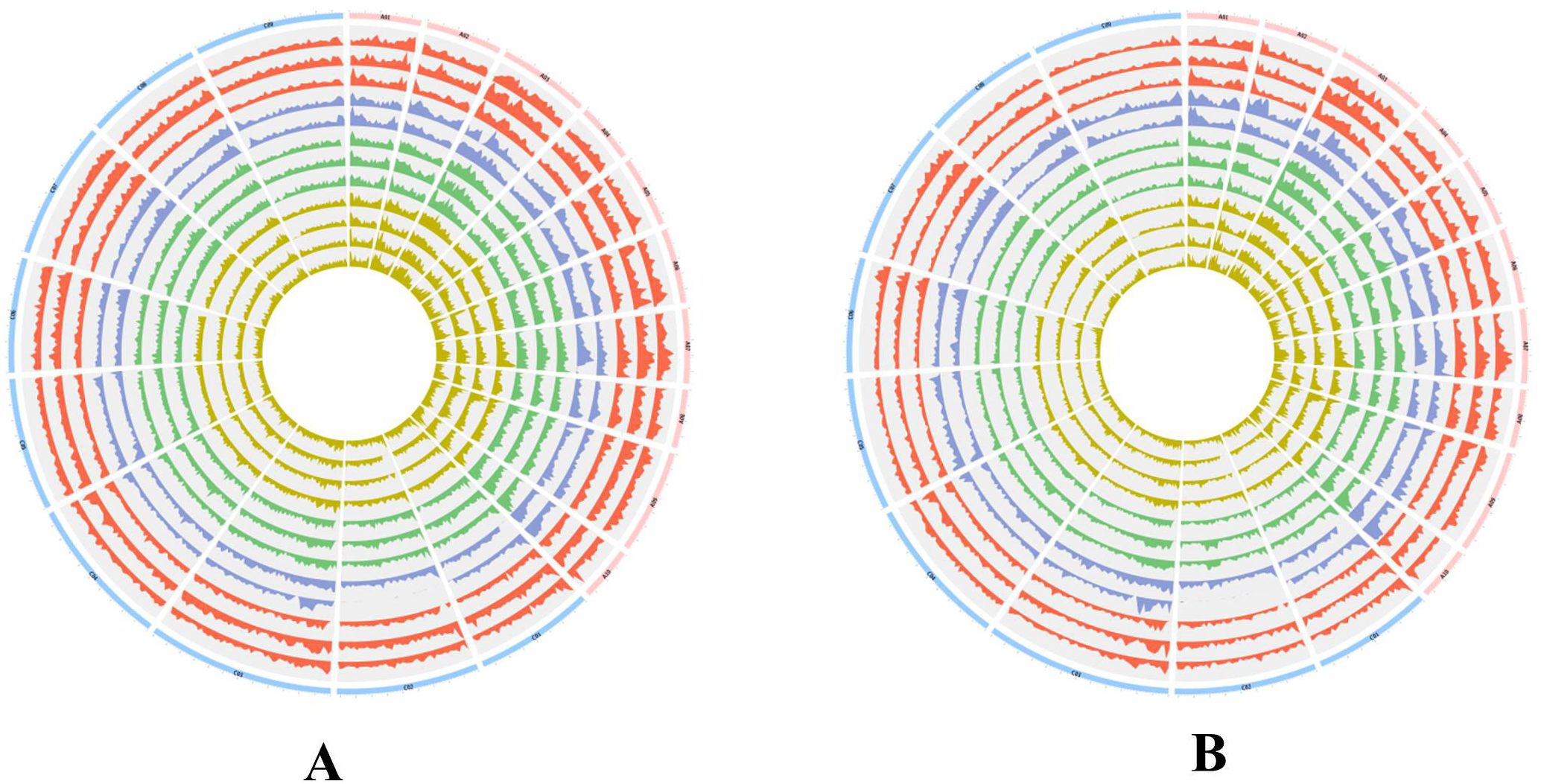
**A.** Circos plot depicting number of small to mid-scale deletion events calculated in 1 Mbp windows across 19 chromosomes of 12 *B. napus* accessions. Each track represents a single genotype in the following order from outside to inside: Express 617, Quinta, Tapidor, R53, No2l27, N99, Westar, PAK85912, Gangan, Shengli, Zheyou7 and ZS11. Colours of tracks represent different types of *B. napus*. The red, blue, green and yellow track colours represent winter-type, synthetic, spring-type and semi-winter accessions, respectively. B. Circos plot depicting the frequency of small to mid-scale insertion events in 1 Mbp windows across 19 chromosomes of 12 *B. napus* genotypes. Each track represents a single genotype in the following order from outside to inside: Express 617, Quinta, Tapidor, R53, No2l27, N99, Westar, PAK85912, Gangan, Shengli, Zheyou7 and ZS11. Colours of tracks represent different types of *B. napus*. The red, blue, green and yellow track colours represent winter-type, synthetic, springtype and semi-winter accessions respectively.

### Implications of long-read sequencing technologies for discovery of functional diversity

Of nine additional SV events we evaluated using PCR, all showed the expected PCR products corresponding to the deletions or insertions predicted by the long-read SV calling. These results underline the apparent effectiveness of long sequence reads for accurately detecting and anchoring insertions/deletions in a broad size range from under 100 bp up to multiple kbp. In contrast, Illumina short reads from regions corresponding to insertions not present in available reference genomes remain un-aligned in alignment-based resequencing approaches, meaning that their genomic localization using short-read data can be achieved only by whole genome *de novo* assembly. Our results in *B. napus* showed that *de novo* SV events appear to occur at an unexpectedly high rate. Hence, it remains unclear how many high-quality reference genomes will be necessary to construct a representative pangenome that captures the majority of the genome-wide functional SV landscape.

This study provides one of the very first insights into genome-wide, gene scale SV linked to important agronomic traits in a major crop species. Recently, Yang et al. (2019) revealed a similar scale of widespread SV by comparing whole-genome assemblies of two diverse maize accessions. However, the cost of genome assembly is still much too high to capture the full extent of species-wide SV in large numbers of genotypes, particularly in species like *B. napus* with dynamic polyploid genomes in which genome rearrangement may even still be ongoing. Our successful verification of 10 out of 10 SV selected events via PCR (Supplementary table S8) gives us high confidence that SV predicted using medium-coverage long-read data with our calling strategy are genuine. This provides a relatively cost-effective method to assay larger germplasm collections without ascertainment bias.

The occurrence of SV events in a size range corresponding to intragenic rearrangements (~100-1000 nt) has been ignored in most crop species in the past, due to the limited resolution of short-read resequencing. Although presence-absence calling from genome-wide SNP array data has been successful in isolated cases in establishing QTL associations (e.g. Gabur et al., 2018a), SNP-based genome-wide association (GWAS) studies are unable to tag causative SV in crops and genome regions in which high levels of LD decay surround the SV events (Zhou et al., 2019). Array-based approaches to call presence absence variations (PAV) or homoeologous exchanges (e.g. Grandke et al. 2016) are therefore likely to ignore potentially functional SV events. Reduced costs, considerably improved read accuracy and significantly increased average read lengths today make long-read sequencing technologies a viable option not only for accurate assemblies of complex plant genomes (Belser et al., 2018), but increasingly also for genome-wide resequencing. Our results suggest that simple reference-based resequencing and alignment with long reads can uncover a new dimension of genetic and genomic diversity associated with important traits in crop plants. Particularly in polyploid plants (Schiessl et al., 2019), this may lead to discovery of previously unknown levels of functional diversity of major interest for breeding and crop adaptation.

## Experimental procedures

### Plant material

We chose 12 *B. napus* genotypes (Table 1) comprising of 3 winter, 4 semi-winter, 3 spring and 2 synthetics (one each of winter and spring).

### DNA isolation for Oxford Nanopore Technology (ONT) sequencing

High molecular weight DNA was isolated using DNA isolation protocol modified from Mayjonade et al. (2016). Young leaves were harvested from rapeseed plants at 4-6 leaf stage and flash frozen using liquid nitrogen. Frozen leaf material was ground to fine powder using a mortar and pestle and transferred to 15 ml Falcon tube. 4-5 ml of pre-heated lysis buffer (1% w/v PVP40, 1% w/v PVP10, 500 mM NaCl, 100mM TRIS pH8, 50 mM EDTA, 1.25% w/v SDS, 1% (w/v) Na_2_S_2_O_5_, 5mM C_4_H_10_O_2_S_2_, 1 % v/v Triton X-100) was added in order to disrupt the cell wall. The lysate was incubated for 30 minutes at 37°C in a thermomixer. 0.3 volumes of 5M Potassium Acetate was added to the lysate and spun at 8000g for 12 minutes at 4°C to precipitates sodium dodecyl sulfate (SDS) and SDS-bound proteins in order to obtain clean DNA. Finally, magnetic beads were used to recover cleaned DNA.

### Library preparation for ONT sequencing

Between 1-3ug of DNA was used to prepare the sequencing library, using the ligation sequencing kit SQK-LSK108 or SQK-LSK109 according to the manufacturer’s recommendations. Genomic DNA was subjected to end repair followed by a bead cleanup. Sequencing adaptors were then ligated to the end-repaired DNA. Finally, the adaptor ligated DNA was once again subjected to bead cleaning. DNA was finally loaded onto an Oxford Nanopore MinION flow cell for sequencing.

### Pacific Biosciences (PacBio) sequencing

Raw PacBio reads originating from 8 genotypes (Quinta, Tapidor, No2127, Westar, Gangan, Shengli, Zheyou7 and ZS11) were downloaded from NCBI short read archive (Accession number PRJNA546246) with the permission from the authors.

## Bioinformatics analysis

### Alignment and SV calling for ONT data

Raw fast5 files obtained by the MinION device were base-called using ONT provided base-caller, Albacore. Raw, uncorrected reads from various flow cells were combined into single fastq file for each genotype. This fastq file was used to align the Nanopore reads to the publically available *B. napus* reference genome assembly Darmor-bzh v4.1 (Chalhoub et al., 2014) and also to a concatenated pseudo-reference assembly comprising the *B. rapa* and *B. oleracea* reference assemblies recently published by Belser et al. (2018), using NGMLR version 0.2.7 (Sedlazeck et al., 2018) with default settings except for “-x ont” flag, representing parameter presets for ONT. NGMLR produced an un-sorted SAM file as an output, which was converted to a sorted BAM file using Samtools version 1.9 (Li et al., 2009). Genomic variants were called using Sniffles version 1.0.10 (Sedlazeck et al., 2018) using the preset parameters.

### Alignment and SV calling for PacBio data

Since 8 PacBio libraries contained nearly 70-80 Gbp of sequencing data, we randomly selected 50 Gbp of data for further analysis in order to obtain quantitatively comparable data to the Nanopore sequencing. This 50 Gbp of data was then aligned as per section 1.4.1 to the publicly available *B. napus* reference and also to the concatenated pseudo-reference assembly, using NGMLR version 0.2.7 with default settings. NGMLR produced an un-sorted SAM file as an output, which was converted to a sorted BAM file using Samtools version 1.9. Genomic variants were called using Sniffles (version 1.0.10) using the preset parameters.

### Quality filtering of the predicted SV events for both ONT and PacBio datasets

We performed a very stringent quality filtering on the sniffles predicted SV events. Since the study was focused on small scale insertions or deletions, we removed all predicted translocations and duplications. Furthermore, it is nearly impossible to validate the authenticity of such SV events, as many may represent mis-positioning of genomic fragments in the reference assembly, we only considered SV scored as “PASS” by Sniffles and ignored those scored as “UNRESOLVED”. Sniffles reports SVs with both within-alignment (AL) and split-read (SR) information. AL-type SV are usually small indels that can be spanned within a single alignment, whereas large or complex events lead to SR alignments (Sedlazeck et al., 2018). To ensure only the high confidence SV were selected, all SV which were not supported by a “within-alignment: AL” flag were discarded. This might lead to an under-estimation and bias in the size distribution of the detectable SV. However, at this point of time the accuracy of publically available genome from *B. napus* is not high enough to distinguish large and complex SV events from assembly errors.

### Calculation of overlap between SV events and the gene models

Quality filtered SV events were overlapped with the gene models from Darmor*-bzh* and also to the combined *B. rapa* and *B. oleracea* reference assemblies using Bedtools intersect (Quinlan and Hall, 2010) using the default parameters. In order to calculate the genome wide frequency of SV events, we also overlapped the quality filtered SV with a bed file containing 1 Mbp windows for the entire genome assembly. The intersect file between the SV events and 1 Mbp windows for the entire genome assembly was then used for plotting the SV distribution along 19 *B. napus* chromosomes, using Circos (Krzywinski et al., 2009). Statistics including length and distribution of quality-filtered SV from the 12 genotypes were calculated with SURVIVOR (Jeffares et al., 2017) and plotted with ggplot2 (Wickham, 2016).

### Construction of a Maximum Likelihood (ML) tree

SV events predicted for each of the 12 genotypes were merged into a single variant calling file (vcf). This combined vcf was then used to force call all the SV events across all 12 genotypes using Sniffles, resulting in a multi-sample vcf. The multi-sample vcf was then converted into PHYLIP format using an in house bash script and used as an input for IQ-TREE version 1.6.12 (Nguyen et al., 2015). The best-fit substitution model for the data was determined by IQ-TREE ModelFinder (Kalyaanamoorthy et al., 2017) and used to construct a phylogenetic tree. The tree was then plotted with FigTree (http://tree.bio.ed.ac.uk/software/figtree/).

### Selection of SV events for PCR validation

We looked at two different agronomically interesting traits in order to prioritize the predicted SV events. Firstly, we analyzed the SV events that might contribute to *Verticillium longisporum* (VL) resistance, using a bi-parental double-haploid population derived from a cross between our sequencing panel genotypes Express 617 and R53. Two QTL were defined for VL resistance on chromosome C01 and C05 by Obermeier et al. (2013). We mainly focused on C05 QTL, as this was described to be the major genetic control for VL resistance. The genetic map used for identifying C05 QTL was based on SSR (Simple Sequence Repeats) and AFLP (Amplified Fragment Length Polymorphism) markers. Therefore, in order to localize the physical position of the QTL on chromosome C05, we anchored the flanking SSR markers (BRMS030_210 and Na12C01_160) to the Darmor-*bzh* version 4.1 assembly and identified a 4.3 Mbp (6,329,426 bp to 10,659,726 bp) region containing 606 genes. 37 and 45 out of the 606 genes were found to contain SV in the form of insertions or deletions in Express 617 and R53 respectively. 17 genes were found to be common among both the genotypes, so were dropped from the prioritized gene set. We further prioritized the candidate genes, if they were annotated as defense response or phenolpropanoid pathway genes. Secondly, we analyzed the SV located within the genes described to be involved in flowering time pathway in *B. napus* as described by Schiessl et al. (2017). Top prioritized SV were then visualized in IGV viewer (Robinson et al., 2017) and selected for PCR validation.

## Authorship

HSC, HTL and RJS conceived the study. HSC, STNA and IAPP generated the Oxford Nanopore long-read sequence data. JS, KL and LG contributed PacBio long-read sequence data. SVS contributed Illumina sequence capture data. HSC, STNA and HTL conducted the experiments and analysed the data. IG, CO, RJS and HTL provided ideas and suggestions for data analysis. HSC and RS drafted the manuscript.

